# CDC50 orthologues in *Plasmodium falciparum* have distinct roles in merozoite egress and trophozoite maturation

**DOI:** 10.1101/2022.01.18.476868

**Authors:** Avnish Patel, Stephanie D. Nofal, Michael J. Blackman, David A. Baker

**Affiliations:** Department of Infection Biology, London School of Hygiene & Tropical Medicine, London, United Kingdom; Malaria Biochemistry Laboratory, The Francis Crick Institute, London, United Kingdom and Department of Infection Biology, London School of Hygiene & Tropical Medicine, London, United Kingdom.

## Abstract

In model organisms P4-ATPases require cell division control protein 50 (CDC50) chaperones for their phospholipid flipping activity. In the malaria parasite, *P. falciparum,* guanylyl cyclase alpha (GCα) is an integral membrane protein that is essential for release (egress) of merozoites from their host erythrocytes. GCα is unusual in that it contains both a C-terminal cyclase domain and an N-terminal P4-ATPase domain of unknown function. We sought to investigate whether any of the three CDC50 orthologues (denoted A, B and C) encoded by *P. falciparum* are required for GCα function. Using gene tagging and conditional gene disruption, we demonstrate that both CDC50B and CDC50C are expressed in the clinically important asexual blood stages and that CDC50B is a binding partner of GCα whereas CDC50C is the binding partner of another putative P4-ATPase, ATP2. Our findings indicate that CDC50B has no essential role for intraerythrocytic parasite maturation but modulates the rate of parasite egress by interacting with GCα for optimal cGMP synthesis. In contrast, CDC50C is essential for blood stage trophozoite maturation. Additionally, we find that the CDC50C-ATP2 complex may influence parasite endocytosis of host cell haemoglobin and consequently hemozoin formation.

## Introduction

*Plasmodium falciparum* is responsible for the majority of malaria mortality and morbidity globally. Whilst there was a sharp reduction in malaria-related deaths between 2000 and 2014 due to increased surveillance, improved control measures and the use of highly effective drug treatments, the decline in cases has halted in recent years. This is thought to be due to the emergence of resistance to insecticides in the *Anopheles* mosquito vector, and to the appearance of parasites resistant to artemisinin combination therapies (ACTs) (1). Given this trend, novel targets must be explored to generate candidates for the drug development pipeline to prevent a future increase in disease burden should the ACTs fail (2).

*P. falciparum* has a complex life cycle, characterised by multiple specialised developmental forms which transition between the mosquito vector and humans (3). Malaria pathology is caused exclusively by the asexual blood stage of the life cycle. Briefly, extracellular merozoites invade host erythrocytes and transform into ring stages for around 24 h. These develop within a membrane-enclosed parasitophorous vacuole to form trophozoites which digest host cell haemoglobin and initiate DNA replication and endomitosis. The resulting schizonts undergo cytokinesis (segmentation) only upon maturation, forming daughter merozoites which are released from the host cell through a highly regulated egress process. The cycle is then re- initiated by the invasion of fresh erythrocytes by the newly released merozoites. A detailed molecular understanding of the biochemical pathways and proteins essential for blood-stage development will inform discovery of novel targeted therapeutics that prevent malaria pathogenesis.

Merozoite egress and invasion are regulated by cyclic nucleotide signalling, conserved elements of which regulate multiple aspects of cell biology in model organisms and across the animal kingdom and can be effectively targeted pharmacologically (4). The second messengers cyclic adenosine monophosphate (cAMP) and cyclic guanosine monophosphate (cGMP), are produced by cyclase enzymes and activate their respective cyclic nucleotide-dependent effector protein kinases PKA and PKG in a concentration-dependent manner (5). The activated kinases phosphorylate downstream targets which carry out effector functions and the cyclic nucleotide signals are then broken down by phosphodiesterases (6, 7). *P. falciparum* uses cyclic nucleotide signalling throughout its complex life cycle (7). Notably, cGMP signalling is required for egress of asexual blood stage merozoites (8, 9) but also egress of gametes (10) and liver stage parasites (11, 12) as well as for ookinete and sporozoite motility (12–14). In contrast, cAMP signalling has been shown to be required for sporozoite apical organelle secretion and invasion of hepatocytes (15), gametocyte deformability (16) and erythrocyte invasion (17).

*P. falciparum* has two guanylyl cyclases. Whilst guanylyl cyclase beta (GCβ) is dispensable in blood stages (9), guanylyl cyclase alpha (GCα) synthesises cGMP in mature blood stage schizonts, where it plays an essential role in activating PKG to trigger egress (18). Both of the malaria parasite GCs are large integral membrane proteins with 22 predicted transmembrane domains (TMDs), the C-terminal segment of which constitutes the paired C1 and C2 guanylyl cyclase catalytic domains. Uniquely for cyclase enzymes, *Plasmodium* GCs (along with apicomplexan and ciliate orthologues) also contain an N-terminal Type IV P-type ATPase (P4- ATPase)-like domain (18–20). In other organisms P4-ATPases transport phospholipids from the outer to the inner leaflet of a lipid bilayer, maintaining lipid asymmetry required for numerous functions including membrane remodeling and vesicle formation (21). Recent studies in the apicomplexan parasite *Toxoplasma* indicate that this domain is critical to the role of its single guanylyl cyclase TgGC in lytic growth, where it is essential for host cell attachment, invasion and motility-dependent egress of tachyzoites (22–25).

In model organisms, P4-ATPases require cell division control protein 50 (CDC50) chaperones for their phospholipid flipping activity (26, 27). CDC50 proteins are integral membrane proteins with two TMDs (28, 29). Studies in yeast have shown that CDC50 binding partners are required for the auto-phosphorylation of the catalytically active aspartic acid residue of the P4-ATPase, which is necessary for completion of the phospholipid flipping reaction cycle (30, 31). *P. falciparum* encodes three putative CDC50 proteins, termed CDC50A (PF3D7_0719500), CDC50B (PF3D7_1133300) and CDC50C (PF3D7_1029400). Previous work in the mouse malaria model *P. yoelii* has shown that CDC50A binds to GCβ and is required for ookinete motility (20). Similarly, TgGC controls egress of tachyzoites (22, 24, 25) and binds to a *Toxoplasma* CDC50 partner which is required for its function (22). However, the functions of *P. falciparum* CDC50 orthologues have not been examined. Here we show that both CDC50B and CDC50C are expressed in the asexual blood stages and that CDC50B interacts with GCα whereas CDC50C is the binding partner of another putative P4-ATPase (ATPase2; PF3D7_1219600). We show that CDC50B modulates the efficiency of parasite egress by interacting with GCα for optimal cGMP synthesis. In contrast, CDC50C is essential for asexual blood stage trophozoite maturation due to a crucial role in endocytosis of host erythrocyte haemoglobin.

## Results

### 1) Generation of genetic tools to investigate the function of the *P. falciparum* CDC50s

To assess the biological functions of CDC50A, B and C in *P. falciparum* blood stages, we generated three transgenic parasite lines designed to allow investigation of subcellular location and the effects of conditional disruption of each CDC50. The transgenics were generated in the genetic background of a 3D7 *P. falciparum* line that stably expresses dimerisable Cre (DiCre), the Cre-recombinase activity of which is induced in the presence of rapamycin (RAP) (32, 33). In each case, the target genes were ‘floxed’ such that treatment with RAP would lead to excision of DNA sequences encoding a C-terminal region containing the second TMD of each protein (Fig 1 A and B); this TMD has been shown in model organism CDC50-ATPase structures to interact with the C-terminal helix of the ATPase binding partner (28, 29). The constructs were designed so that following homologous recombination, the genes were also modified by fusion to sequences encoding a C-terminal triple hemagglutinin (HA) epitope tag.

**Figure 1.**
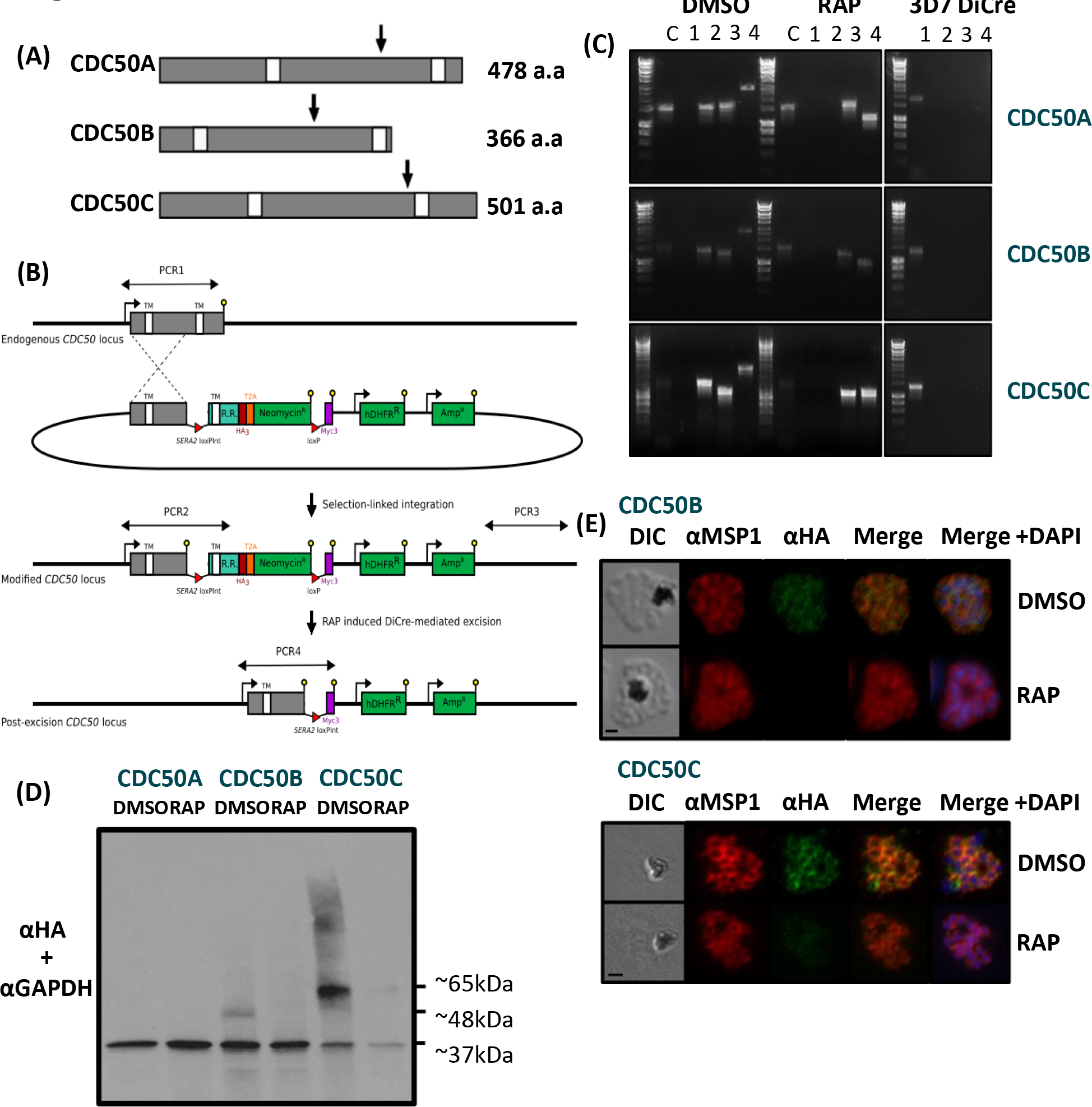
(1A) Representation of the three CDC50 proteins in *P. falciparum* displayed from N- to C- terminus in a relative scale. White boxes indicate transmembrane helices (TMDs) as predicted by the TMHMM server (59). Arrows indicate the point at which the protein products are truncated when the corresponding modified locus is excised in transgenically-modified parasite lines. For CDC50A this is from Phe341, for CDC50B His235 and CDC50C from Glu383. (1B) Schematic representation of the SLI strategy (58) used to produce the three CDC50 DiCre lines and resultant RAP-induced disruption of the modified genes. Double-headed arrows represent the regions amplified by PCR in (1C). Red arrowheads represent *loxP* sites, yellow lollipops represent translational stop codons, white boxes indicate TMDs and light blue boxes indicate regions of re-codonised sequence (R.R.). (1C) Diagnostic PCR analysis of gDNA from transgenic CDC50 parasite lines verifying successful modification of target loci by SLI to produce CDC50A-HA:loxP, CDC50B-HA:loxP and CDC50C- HA:loxP. Efficient excision of ’floxed’ sequences is observed upon treatment with RAP for all lines. Track C represents amplification of a control locus (PKAc) to check gDNA integrity. PCRs 1- 4 are represented in the schematic locus in panel 1B. PCR 1 screens for the WT locus, PCR 2 for 5’ integration, PCR 3 for 3’ integration and PCR 4 for the excision of the ’floxed’ sequence. See Table 1 for sequences of all primers used for PCR. Sizes for expected amplification products are as follows: C, control locus (primers 16 and 17) 1642b.p. CDC50A; PCR 1 (primers 21 and 22) 1842b.p, PCR2 (primers 21 and 18) 1613b.p, PCR3 (primers 20 and 22) 1670b.p and PCR 4 (primers 21 and 19) 2863b.p (DMSO), 1169b.p (RAP). CDC50B; PCR 1 (primers 23 and 24) 1423b.p, PCR2 (primers 23 and 18) 1457b.p, PCR3 (primers 20 and 24) 1321b.p and PCR 4 (primers 23 and 19) 2707b.p (DMSO), 1010b.p (RAP). CDC50C; PCR 1 (primers 25 and 26) 1369b.p, PCR2 (primers 25 and 18) 1602b.p, PCR3 (primers 20 and 26) 1172b.p and PCR 4 (primers 25 and 19) 2852b.p (DMSO), 1369b.p (RAP). (1D) Western blot analysis of expression (DMSO) and ablation (RAP) of CDC50A-HA, CDC50B-HA and CDC50C-HA from highly synchronous late stage schizonts in the respective transgenic parasite lines. Expression of GAPDH (PF3D7_1462800) is shown as a loading control. No expression of CDC50A-HA was detected. Predicted molecular weights of CDC50B-HA, CDC50C- HA and GAPDH are indicated. (1E) IFA analysis showing the diffuse peripheral localisation of CDC50B-HA and CDCD50C-HA and the loss of expression upon RAP treatment. Over 99% of all RAP-treated CDC50B-HA:loxP and CDC50C-HA:loxP schizonts examined by IFA were diminished in HA expression in three independent experiments. Scale bar, 2 µm.

**Table 1.**
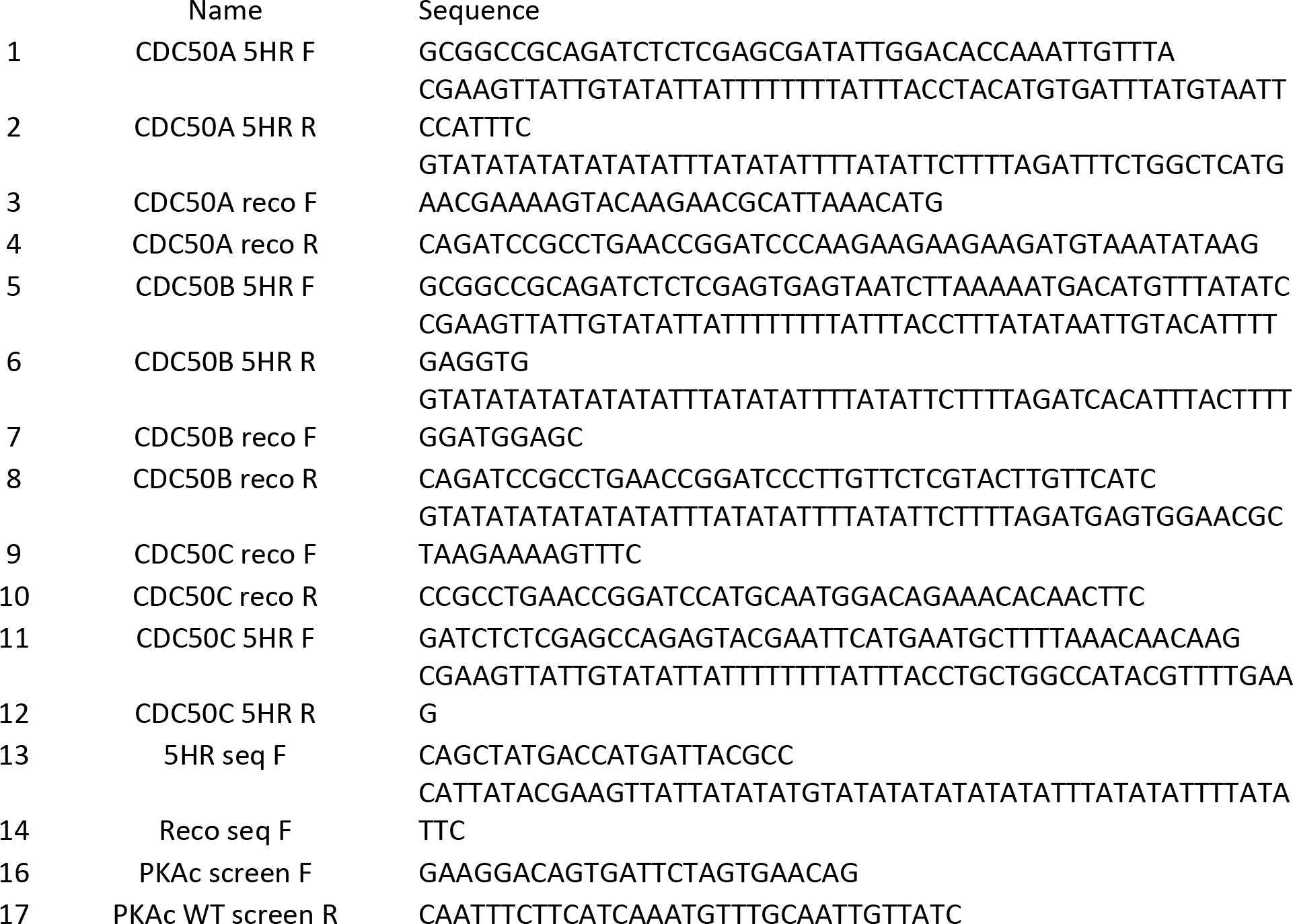
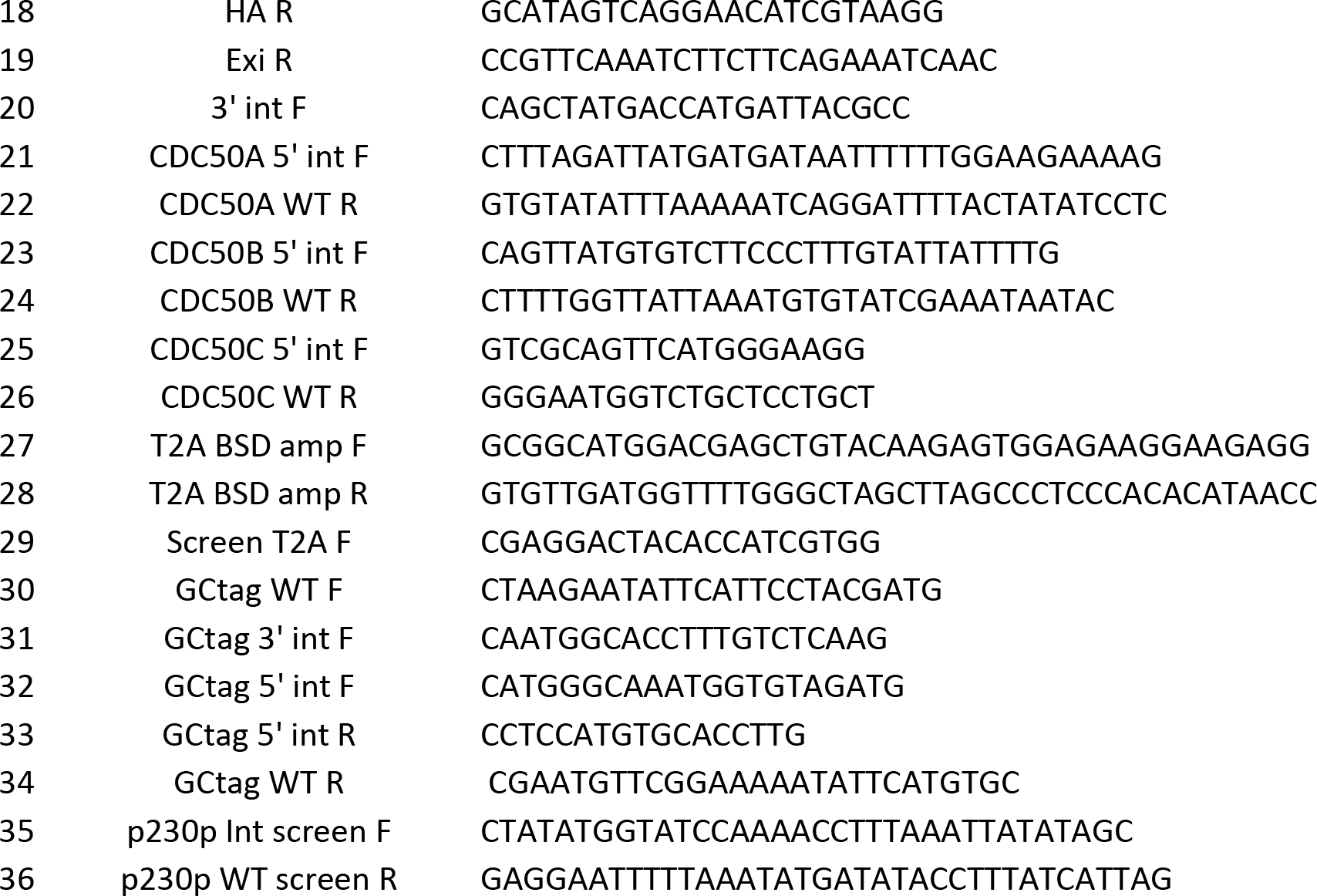
oligonucleotide primers used in this study

Successful modification of the target genes was verified by PCR (Fig 1C), and expression and RAP-induced excision of tagged CDC50A-HA, CDC50B-HA and CDC50C-HA fragments in the respective transgenic parasites (termed CDC50A-HA:loxP, CDC50C-HA:loxP and CDC50B- HA:loxP) was confirmed by western blot (Fig 1D). Immunofluorescence analysis (IFA) of the transgenic lines (Fig 1E) revealed a diffuse, partly peripheral signal in individual merozoites within mature segmented schizonts for both CDC50B-HA and CDC50C-HA. This was similar to the pattern observed upon co-staining with the plasma membrane marker, merozoite surface protein 1 (MSP1). Whilst successful tagging and floxing of the *CDC50A* gene was also confirmed by PCR and Sanger sequencing, no protein expression could be detected in asexual blood stages. This suggested that CDC50A is not expressed in asexual stages, consistent with findings in *P. yoelii* where it is expressed only in gametocyte and mosquito stages (20). Alternatively, since *P. falciparum* transcriptomic data indicates that the *CDC50A* gene is transcribed in schizont stages (34), the protein may be expressed but rapidly degraded as its GCβ binding partner is not present in asexual blood stages (9, 19).

### 2) Of the three isoforms, only CDC50C is essential for blood stage parasite growth

To investigate the essentiality of CDC50A, CDC50B and CDC50C, highly synchronised ring-stage cultures of each DiCre transgenic line were treated with RAP to induce excision of the sequence encoding the C-terminal TMD of each protein (Fig 1C), and parasite replication assessed using flow cytometry (Fig 3A). RAP-treated CDC50A-HA:loxP and CDC50B-HA:loxP parasites displayed no significant growth inhibition over three cycles compared to matched control (DMSO-treated) parasites (Fig 2A). In contrast, CDC50C-HA:loxP parasites underwent complete growth arrest after cycle 1. Examination of the parasites by Giemsa-staining showed that whilst new rings went on to form schizonts in DMSO-treated WT parasites, RAP-treated CDC50C-HA:loxP rings did not develop beyond the early trophozoite stage and eventually collapsed into small vacuoles (Fig 2B). It was concluded that CDC50C is essential for asexual blood stage survival.

**Figure 2.**
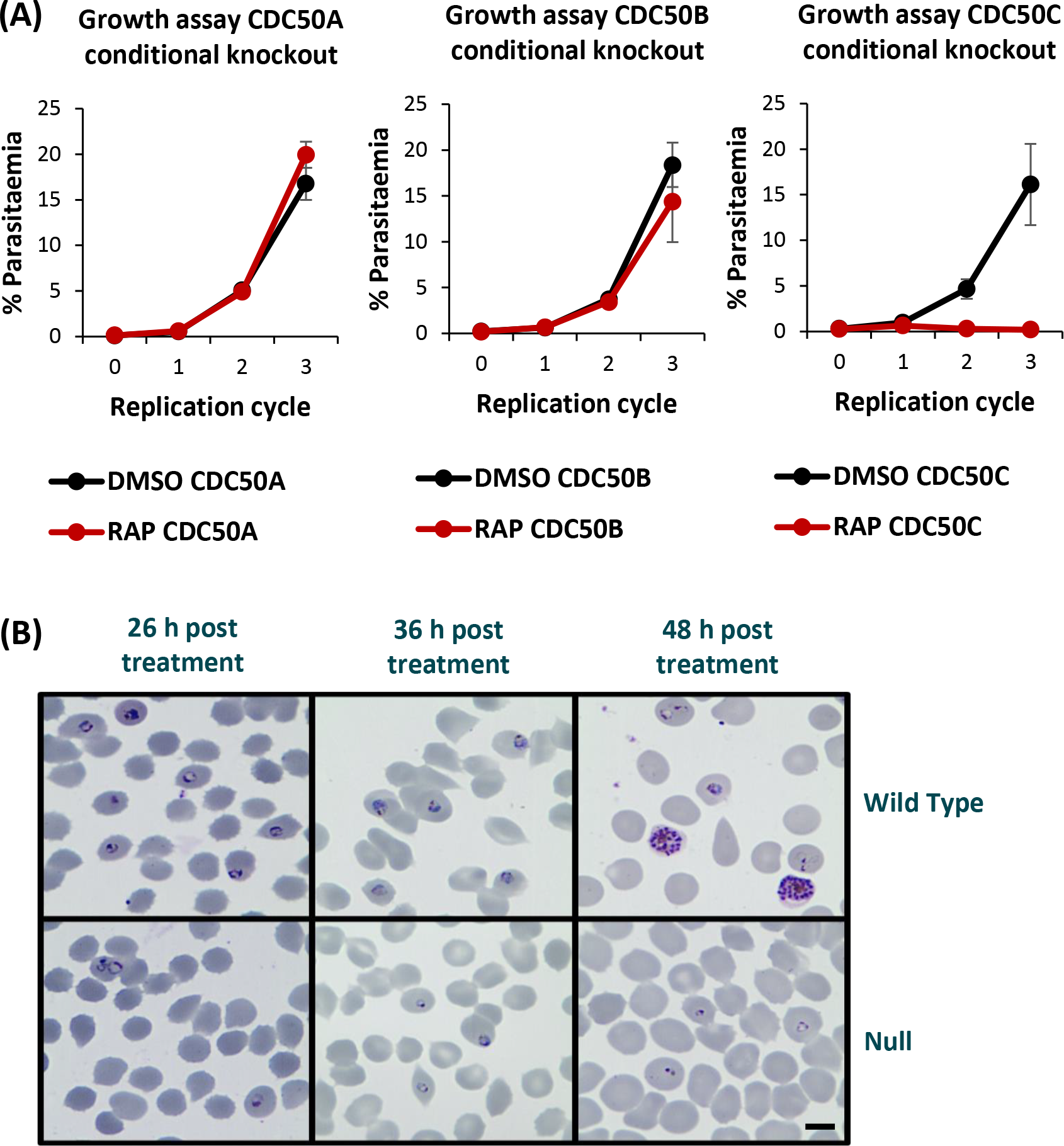
(2A) Growth curves showing parasitaemia as measured by flow cytometry of CDC50A-HA:loxP, CDC50B-HA:loxP and CDC50C-HA:loxP parasites treated with DMSO (vehicle only control) or RAP. Means from 3 biological replicates are plotted. Error bars, SD. (2B) Giemsa-stained thin blood films showing ring-stage parasites following egress of synchronous DMSO and RAP-treated CDC50C-HA:loxP schizonts. Ring formation occurs in RAP- treated CDC50C-HA parasites, but the parasites did not develop beyond the early trophozoite stage and eventually collapsed into small vacuoles.

**Figure 3.**
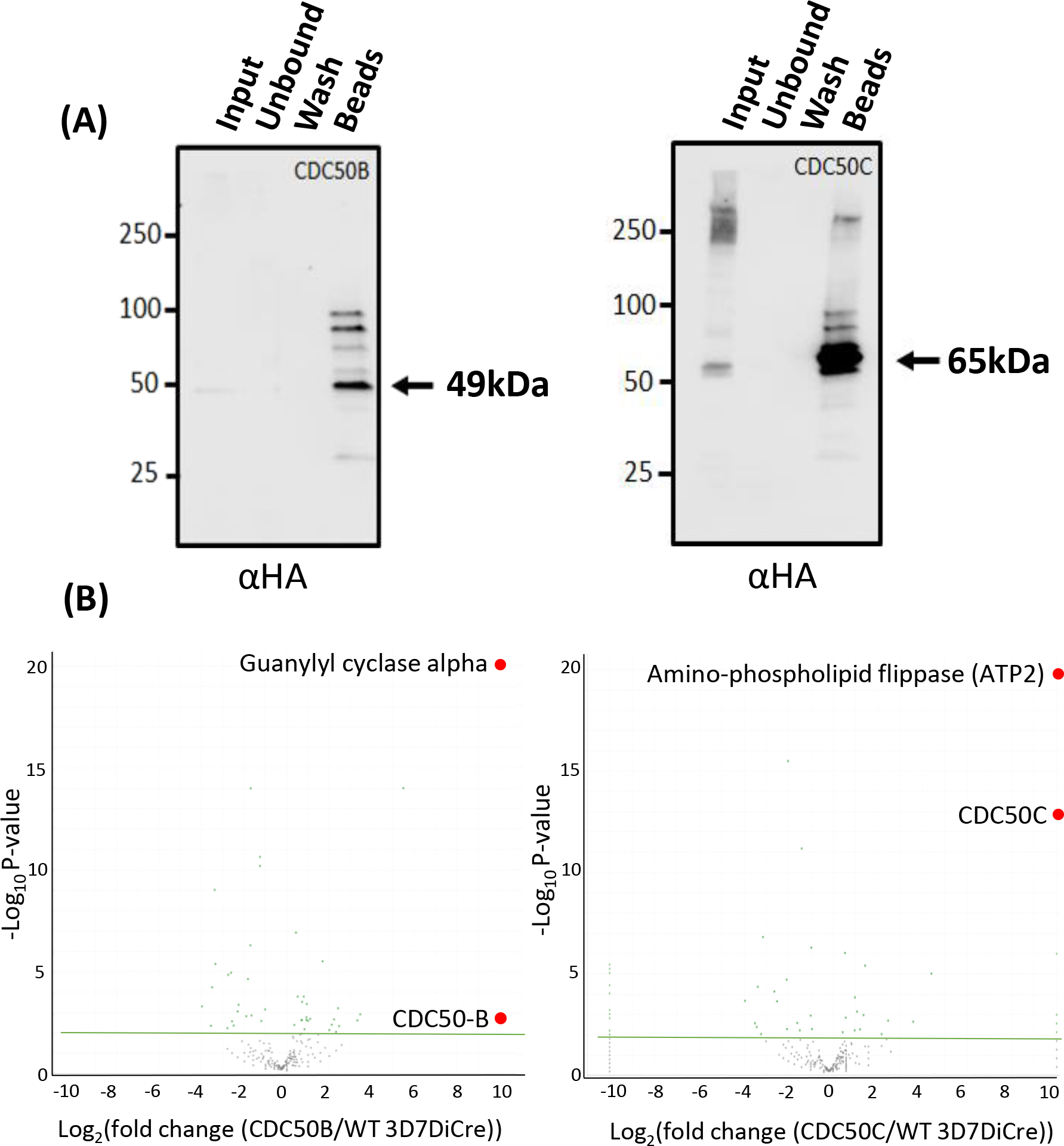
(3A) Western blots demonstrate efficient immunoprecipitation (IP) of CDC50B-HA and CDC50C- HA from schizont extracts. Large black arrows indicate the predicted mass of each protein. Images are representative of 2 biological repeats. (3B) Mass spectrometric identification of interacting partners of CDC50B and CDC50C. Volcano plot of P values versus the corresponding log2 fold change in abundance compared to 3D7DiCre control samples (Fischer’s exact test). Plotted by analysing proteins enriched through IP (panel 3A) by mass spectrometry. Green line indicates p=-2log10 and green dots represent peptides where p<-2log10. Peptides for GCα and ATP2 were enriched to p<-19log10.

### 3) CDC50B and CDC50C bind to distinct parasite flippase partners

In other organisms, CDC50 proteins interact with their cognate P4-ATPases and are required for their activity (28–31). To determine whether CDC50B and CDC50C interact with P4-ATPases in *P. falciparum* blood stage development we performed immuno-precipitation (IP) experiments from extracts of highly synchronised CDC50B-HA:loxP and CDC50C-HA:loxP schizonts. Western blot analysis confirmed the expected enrichment of the HA-tagged proteins from schizont lysates (Fig 3A). The immuno-precipitated material was then analysed by mass spectrometry in comparison with mock IP samples to confirm this and identify co-precipitating protein species. This confirmed high levels of enrichment of the HA-tagged CDC50 bait proteins (Fig 3B). In addition, in the case of the CDC50B experiments we detected a >9 log2 enrichment of peptides derived from GCα (Fig 3B), whilst in the CDC50C IPs we detected >9 log2 enrichment of peptides mapping to another putative P4-ATPase, a putative amino-phospholipid flippase PF3D7_1219600 (ATP2) (Fig3B). No other proteins were as significantly enriched in each pull-down. These results strongly suggest that CDC50B is a co-factor for GCα and CDC50C is a co- factor for ATP2.

### 4) CDC50B is n ot r equ i r ed f or G Cα exp r essi on or tr af fi cki n g, b ut is crucial for optimal cGMP synthesis required for egress

GCα has a key role in egress, as the source of cGMP required for PKG activation (18). Having determined that CDC50B interacts with GCα, we next investigated whether CDC50B also has a role in parasite egress.

To do this, we first compared the egress kinetics of mature DMSO- and RAP-treated CDC50B- HA:loxP schizonts by monitoring the appearance in culture supernatants over time of proteolytically processed forms of the PV protein serine repeat antigen 5 (SERA5), as a proxy for egress (35). As shown in Fig 4A, this revealed a marked reduction in the rate of egress over the sampling period in RAP-treated CDC50B-HA:loxP parasites as compared to control DMSO-treated counterparts. Densitometric quantitation of data from three independent biological replicate experiments indicated that CDC50B null schizonts undergo ∼50% less egress than WT controls when sampled over two hours (Fig S1). This was not due to a delay in schizont development, since Giemsa staining of DMSO- and RAP-treated CDC50B schizonts showed no detectable delay in parasite maturation, and analysis of DNA content by flow cytometry indicated no significant differences between formation of DMSO- and RAP-treated CDC50B schizonts (Fig 4A lower).

**Figure 4.**
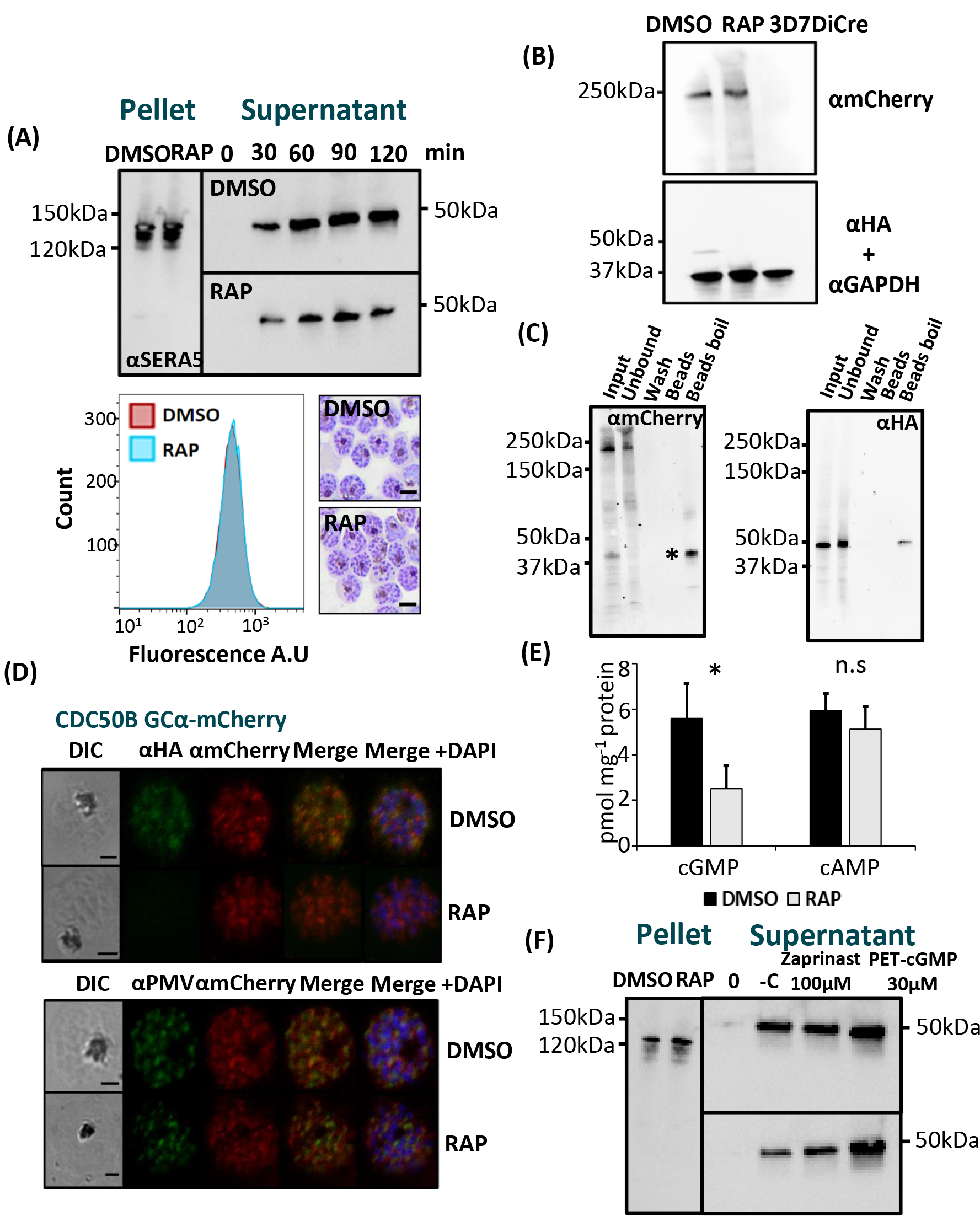
(4A) Western blot analysis monitoring egress kinetics of DMSO- and RAP-treated CDC50B- HA:loxP schizonts. The diminished detection of the SERA5 p50 proteolytic fragment in culture supernatants of RAP-treated CDC50B-HA:loxP parasites indicates an impaired egress rate in the absence of CDC50B-HA. Lower panel left, histograms of DNA (SYBR green) staining of CDC50B DMSO- or RAP-treated schizonts. 10,000 cells were counted per treatment. Image representative of 3 biological repeats. Lower panel right, Giemsa-stained thin blood films of Percoll-purified CDC50B DMSO-and RAP-treated schizonts. No delay in schizont maturation is evident in RAP-treated parasites. Images representative of 3 biological repeats Scale bar, 5µm. (4B) Western blot analysis of DMSO- and RAP-treated CDC50B-HA:loxP GCα-mCherry and control 3D7DiCre schizonts. The top panel shows a ∼250 kDa fragment detected by an mCherry antibody that is absent in control (untagged) schizont lysates. The lower panel shows the same samples probed with an anti-HA antibody and an anti-GAPDH (PF3D7_1462800) loading control antibody. (4C) Immuno-precipitation of GCα-mCherry. Samples were loaded in duplicate and probed for mCherry (left panel) and HA epitope (left panel). * denotes the ∼40 kDa degradation product of GCα-mCherry observed after enrichment and boiling of the RFP-trap beads, suggesting that GCα was prone to proteolysis under the conditions used to promote binding or degrades when heated. (4D) IFA showing the localisation of CDC50B and GCα mCherry in RAP and DMSO treated schizonts, top panel. Bottom panel, localisation of GCα mCherry and PMV (PF3D7_1323500), an ER marker in RAP- and DMSO-treated schizonts. Scale bar, 2 µm. (4E) Quantification of cyclic nucleotide levels in tightly synchronised DMSO- and RAP-treated CDC50B mature schizonts by direct ELISA. Means are shown from 3 biological repeats plotted, error bars, SD. n.s = not significant, * P<0.05, Student’s t test. (4F) Restoration of egress of RAP-treated CDC50B schizonts by treatment with zaprinast or PET- cGMP. Supernatant and pellet samples were taken at time point 0 post washing with RPMI to control for parasite numbers and egress. Samples were then taken at 60 min post incubation at 37 °C. -C represents no treatment. Image is representative of 3 biological repeats.

In *T. gondii*, the orthologue of CDC50B is required for correct subcellular trafficking of TgGC (22). To investigate whether this is also true in *P. falciparum*, we used a CRISPR-Cas9-based approach to fuse GCα to a C-terminal mCherry tag in the CDC50B-HA:loxP line, creating a parasite line called CDC50B-HA:loxP GCα-mCherry (Fig S2). We failed to detect the tagged protein directly by fluorescence microscopy, possibly due to the previously reported very low abundance of GCα (18). However, western blot revealed a ∼250 kDa signal in schizont lysates from CDC50B-HA:loxP GCα-mCherry schizonts, likely representing a proteolytic fragment of the tagged protein, since GCα is prone to proteolytic degradation in both *P. falciparum* and *T. gondi* (18, 22) (Fig 4B). Interestingly, western blot of extracts of RAP-treated CDC50B-HA:loxP GCα- mCherry schizonts indicated that there was no marked reduction in the levels of GCα in the absence of CDC50B (Fig 4B). Exploiting the tagged GCα-mCherry line, we sought to confirm whether CDC50B was co-precipitated when GCα-mCherry was immuno-precipitated using RFP- trap beads. Co-precipitation of CDC50B was observed, confirming CDC50B binding by GCα- mCherry (Fig 4C).

To examine the role of CDC50B in trafficking of GCα, we used an anti-mCherry antibody to localize GCα-mCherry by IFA in RAP-treated CDC50B-HA:loxP GCα-mCherry parasites. This revealed no obvious mis-localisation of GCα in the absence of CDC50B, with a similar, diffuse signal detectable in both RAP- and DMSO-treated schizonts. In addition, in contrast with *T. gondii* (20), no mislocalisation of GCα in the ER or secretory pathway in the absence of CDC50B was detected, as judged by co-localisation with the ER marker plasmepsin V (PMV) (Fig 4E).

Taken together, these results indicate that CDC50B binding is not important for the correct trafficking or stable expression of GCα. To seek more insight into the egress defect, we investigated whether ablation of CDC50B resulted in changes in cyclic nucleotide levels. To do this, we assayed extracts of DMSO- and RAP-treated CDC50B-HA:loxP schizonts by ELISA to quantitate cGMP and cAMP levels. This showed that CDC50B null parasites contained 53.67% (±12.16%) less cGMP than DMSO-treated controls, whilst no significant difference in cAMP levels was observed (Fig 4F). These reduced cGMP levels suggested that binding of CDC50B to GCα might be required for maximal GCα cyclase activity. To test this, we investigated whether the defect in egress of CDC50B null schizonts could be reversed by the addition of compounds that stimulate or mimic elevated cGMP levels. Egress was monitored in the presence and absence of the PDE inhibitor zaprinast or PET-cGMP, a membrane-permeable cGMP analogue known to activate parasite PKG (18, 36). Treatment with either compound restored egress of CDC50B null schizonts to levels similar to those observed in control CDC50B-HA:loxP schizonts, confirming the requirement of CDC50B for optimal cGMP synthesis by GCα (Fig 4G).

In view of the above results, as well as the previous observation that a lipid co-factor may stimulate egress in *Toxoplasma* and *P. falciparum* (22, 37), we examined whether the potential phospholipid flippase activity of the P4-ATPase domain of GCα might be modulated by CDC50B binding. To do this, we investigated whether uptake of fluorescently-labelled PS, PE and PC were affected following excision of CDC50B. DMSO- or RAP-treated CDC50B-HA:loxP late schizonts were incubated with fluorescent lipids, then analysed by flow-cytometry to determine their ability to accumulate lipids. We found no significant difference in the bulk uptake of measured lipids in schizonts in the presence or absence of CDC50B (Fig S3).

### 5) CDC50C is not required for lipid uptake or protein export, but plays an essential role in haemoglobin uptake from the host erythrocyte

As described above, RAP treatment of synchronous, newly invaded CDC50C-HA:loxP rings produced a CDC50C null parasite population that displayed a trophozoite arrest phenotype (Fig 2B), suggesting an essential role for CDC50C in the trophozoite-to-schizont transition. Given this evidence that CDC50C plays a very different role to that of CDC50B, we decided to interrogate more precisely the functional role of CDC50C. Previous transcriptomic analysis has shown that CDC50C is transcribed throughout the asexual blood stage cycle, with relatively low levels of transcription in rings increasing to a peak in mature schizont stages (34). Consistent with this transcriptional profile, immuno-staining of CDC50C-HA:loxP parasites detected expression in ring, trophozoite and schizont stages (Fig S4). Co-staining of CDC50C in trophozoites with antibodies to ERD2 (a Golgi marker), PMV (an ER marker) or EXP2 (a PVM marker) showed that CDC50C displayed a diffuse cytosolic staining with no clear subcellular localisation (Fig5A).

Phospholipid flippases have been shown to contribute directly to cellular lipid uptake (38–40). Initially we speculated that the growth arrest of CDC50C null trophozoites may be due to a dysregulation of lipid uptake as a result of loss of function of the putative amino-phospholipid flippase ATP2 partner. To test this notion, we labelled live RAP- and DMSO-treated CDC50C- HA:loxP trophozoites with the fluorescent amino-phospholipid analogues NBD-PC, NBD-PE and NBD-PS. No discernable difference in lipid uptake between CDC50C null and WT trophozoites was observed by microscopy (Fig 5B), suggesting that CDC50C plays no essential role in uptake of these phospholipids.

**Figure 5.**
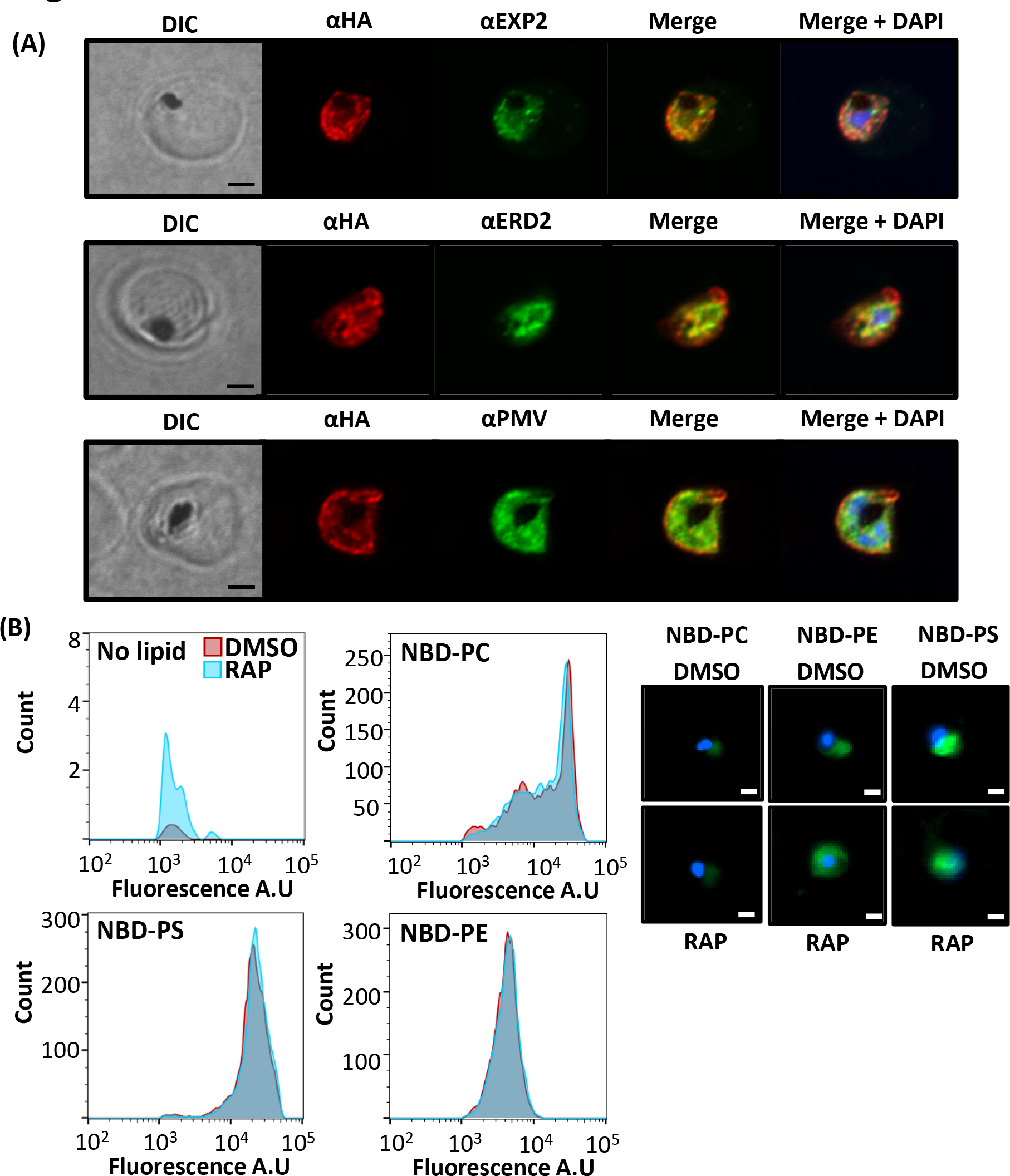
(5A) Airyscan confocal analysis of IFAs of CDC50-C trophozoites co-localised with EXP2 (PF3D7_1471100) an exported PV protein, ERD2 a Golgi marker (PF3D7_1353600) and PMV an ER marker (PF3D7_1323500). Scale bar, 2 µm. (5B) Flow cytometry analysis of fluorescent lipid uptake in live WT (DMSO) and CDC50C null (RAP) trophozoites labelled at 36 h post invasion. Histograms are overlayed each representing 10,000 cells for each treatment. Cells were gated for DNA content and further for green fluorescence. No detectable shift in histogram curves was seen for each lipid in RAP-treated samples. Data are representative of one of three biological repeats, each of which showed the same outcome. Control samples, with no lipid added, were analysed to validate the gating protocol for lipid signal. Right panel shows examples of the stained cells visualised by fluorescence microscopy. Scale bar 2 µm.

In model organisms, flippases also contribute to the production and maintenance of membrane asymmetry required for generation of trafficking vesicles, with specific flippases influencing exocytosis or endocytosis pathways (41–44). Given that lipid uptake was unaffected in the absence of CDC50C, we considered it plausible that the ATP2-CDC50C complex may contribute to lipid homeostasis and trafficking in an analogous manner. *P. falciparum* trophozoites re- model their intracellular environment to create new permeation pathways that enable export of a wide variety of proteins into the host RBC via exocytosis, a process which is essential for trophozoite development (45). To examine whether trophozoite death in CDC50C null parasites could be attributed to changes in protein exocytosis, control or RAP-treated CDC50C-HA:loxP ring stage parasites were allowed to develop into trophozoites then analysed by IFA to determine the localisation of skeleton binding protein (SBP), a prominent exported protein in trophozoites. Puncta of SBP, characteristic of export, were evident within the RBC cytosol in both DMSO- and RAP-treated CDC50C-HA:loxP trophozoites (Fig 6A), and quantification of these puncta indicated that there was no significant change in the levels of export of SBP in CDC50C null trophozoites (Fig 6B). This suggested that loss of CDC50C has no impact on bulk protein export during trophozoite development.

**Figure 6.**
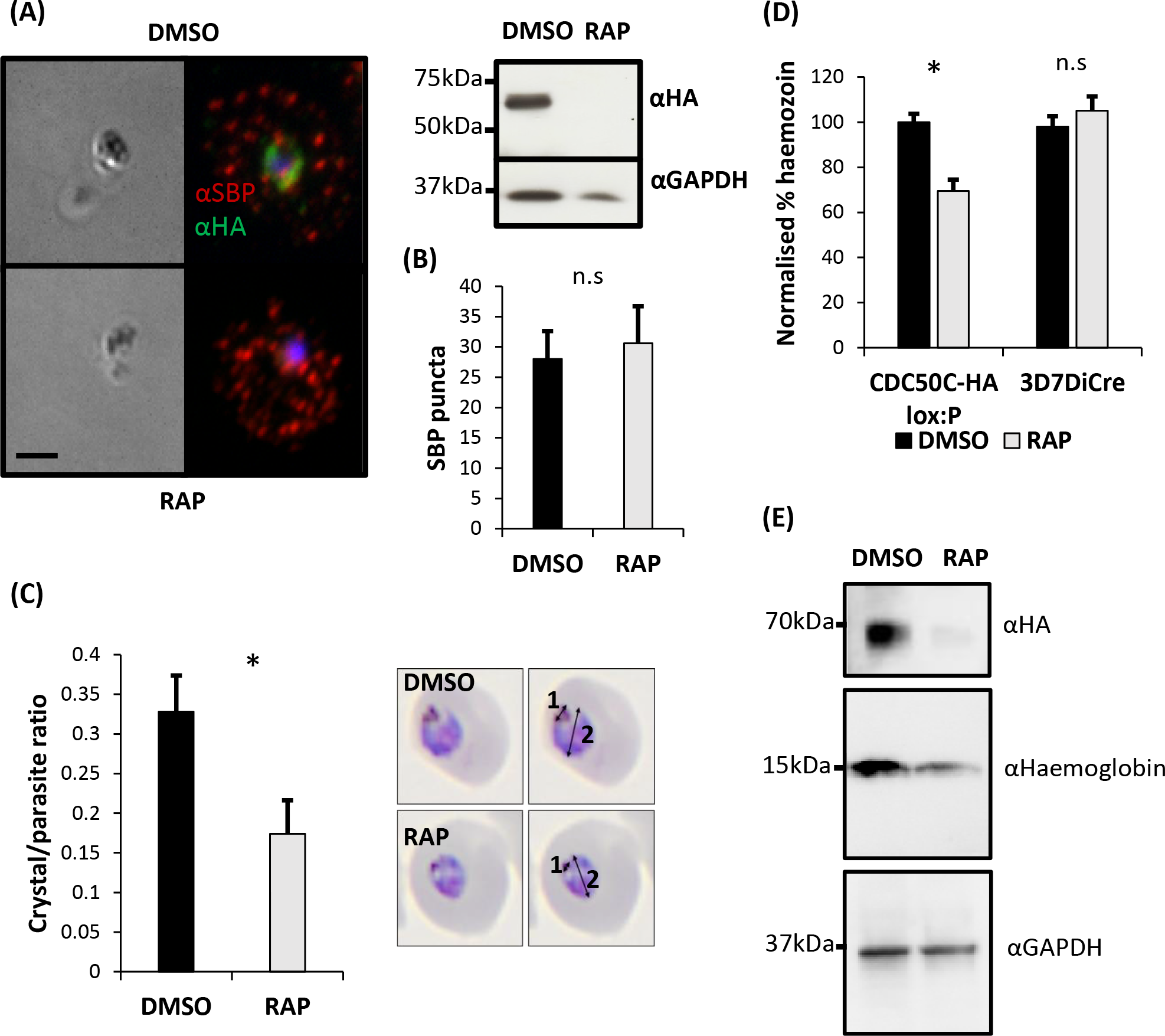
(6A) IFA imaging of DMSO- and RAP-treated CDC50C-HA:loxP trophozoites fixed at 36 h post invasion indicates no defect in export of skeleton binding protein (SBP). Right, western blot showing absence of CDC50C-HA in RAP-treated trophozoites. Scale bar, 2 µm. (6B) Quantification of SBP puncta in DMSO- and RAP-treated CDC50C trophozoites. 66 parasites were counted from 2 individual biological replicates for SBP puncta in ImageJ. Mean values are plotted. Error bars, SD. n.s = not significant, Student’s t test. (6C) CDC50 null parasites produce smaller hemozoin crystals. Thin blood films were made from tightly synchronized DMSO- and RAP-treated CDC50C at 36 h post invasion. Inset, length of the hemozoin crystal (measurement 1) and parasite (measurement 2) were performed in ImageJ on imaged Giemsa-stained smears of DMSO- and RAP-treated CDC50C trophozoites. A ratio of crystal length to parasite length was derived by dividing measurement 1 by measurement 2. In total 46 parasites were measured from 3 independent biological repeats of CDC50B excision or control treatment. Mean values are plotted. Error bars, SD. * P<0.05, Student’s t test. (6D) Spectrophometric quantification of hemozoin content of CDC50C WT (DMSO) and null (RAP) trophozoites. Highly synchronised CDC50C ring stage cultures were treated with DMSO (control) or RAP. Cultures were then harvested at 36 h post invasion. Hemozoin was purified using established methods (46) and quantified by absorbance at 410 nm. Means are plotted for 3 independent biological repeats. Error bars, SD. n.s = not significant, * P<0.05, Student’s t test. (6E) Western blot analysis of the haemoglobin content of saponin lysed CDC50-C WT and null trophozoites. Highly synchronised CDC50C ring stage cultures were treated with DMSO control or RAP. Cultures were then harvested at 36 h post invasion. Lysates were probed for the presence of CDC50-C by HA staining. Haemoglobin content was probed alongside GAPDH as a loading control. Data representative of 3 biological repeats.

As protein export and possibly exocytosis of CDC50C null trophozoites was unaffected we lastly examined an essential endocytotic process. As intracellular asexual blood stage malaria parasites develop, they endocytose and digest host erythrocyte haemoglobin. A major by- product of this catabolic process is the sequestration of haem in the form of a characteristic crystalline product called haemozoin, which accumulates in the parasite digestive vacuole as large, refractile complexes that are easily visible by light microscopy. Microscopic examination of Giemsa-stained thin blood films indicated that whilst CDC50C null trophozoites displayed an apparently normal morphology, the hemozoin crystals appeared smaller than those of DMSO-treated controls (Fig 6C right). To examine this in greater detail, we compared the ratio of the hemozoin crystal length to that of the parasite length in RAP- and DMSO-treated CDC50C null trophozoites. This confirmed a significantly decreased ratio in CDC50C null trophozoites (Fig 6C), suggesting a CDC50C-dependent defect in haemozoin formation. To further examine this, we purified and quantified haemozoin from parallel populations of RAP- and DMSO-treated CDC50C-HA:loxP trophozoites (46). As shown in Fig 6D, this revealed ∼30% less hemozoin in trophozoites lacking CDC50C. To investigate whether uptake of haemoglobin was affected in CDC50C null trophozoites, RAP- and DMSO-treated CDC50C-HA:loxP trophozoites were harvested at 36 h post invasion, released from their host cells using saponin, and levels of intra-parasite haemoglobin quantified by western blotting (Fig 6E). The results indicated that the levels of haemoglobin within CDC50C null trophozoites was significantly reduced compared with control counterparts (Fig S5). Collectively, these data support a role for CDC50C in uptake and digestion of host erythrocyte haemoglobin that is essential for parasite development.

## Discussion

In this study we have shown that CDC50B and CDCD50C proteins are expressed during asexual blood stage development, and that they each bind to different putative P4-ATPase flippases - GCα and ATP2 respectively - which function at different developmental stages of the asexual blood stage cycle. CDC50B is dispensable for intraerythrocytic development, as is its orthologue in *P. yoelii* (20). However, we find that cGMP levels are reduced in CDC50B null parasites as well as rates of egress. This defect can be rescued by treatment of CDC50B null parasites with either the PDE inhibitor zaprinast or PET-cGMP, membrane-permeable cGMP analogue known to be capable of activating apicomplexan PKGs (18). As no growth defect was observed in CDC50B null parasites, it seems that whilst egress rates are reduced, CDC50B parasites are egress competent and do eventually egress. Collectively, our results strongly suggest that CDC50B acts to enhance cGMP synthesis by GCα. Recent studies in *T. gondii* have shown that the single GC (TgGC) also binds to a CDC50 designated CDC50.1 (22). In contrast to the present study, knockdown of CDC50.1 resulted in mis-localisation of TgGC and a block in egress of *T. gondii* tachyzoites (22). The egress block could be rescued by adding a PDE inhibitor, implying that in the absence of CDC50.1, TgGC remains functional but produces cGMP with reduced efficiency (20). We observed no detectable mislocalisation of *P. falciparum* GCα in the absence of CDC50B, with normal growth of parasites albeit with reduced egress rates, indicating differences between the genera. We speculate that this may be due to differences in the threshold levels of cGMP required to activate PKG to trigger egress in each species.

Importantly, our work adds to the evidence supporting a role for CDC50s and the P4-ATPase domain of apicomplexan GCs acting functionally to stimulate maximal cGMP production required for egress. By analogy with other CDC50-flippase interactions, we can deduce that this occurs through CDC50B binding to the P4-ATPase domain of GCα. Modulation of the activity of the C-terminal cyclase domain by the P4-ATPase may integrate a lipid-mediated trigger for egress, potentially by phosphatidic acid as shown in *T. gondii* (22) or phosphatidylcholine (PC) as recently indicated in *P. falciparum* (37). However, we did not observe changes in the bulk uptake of fluorescent PC by schizonts lacking CDC50B. Recent structural examination of a human P4-ATPase:CDC50 complex has shown that CDC50 forms an intimate interaction with the TMDs of the P4-ATPase partner, with the loop domain between the two TMDs of the CDC50 forming an anti-parallel beta-sheet structure that contacts the luminal side of the transmembrane loops of the P4-ATPase. Human CDC50 is glycosylated at several conserved asparagine residues, and the structure showed that interactions between CDC50 glycan moieties and P4-ATPase stabilise the functional complex (29). Intriguingly an alignment of human CDC50a and the three *P. falciparum* CDC50s indicates that Asn180, at which glycosylation has been shown to interact structurally with its partner P4-ATPase (29), is absent from CDC50B but conserved in both CDC50A and C (Fig S6). N-glycosylation has been observed in *P. falciparum* (47), but this finding raises the possibility that CDC50B may be non-glycosylated. This observation may explain the finding in *P. yoelii* that GCβ is degraded in the absence of its partner CDC50, suggesting that GCβ is highly reliant on interactions with its CDC50 partner (CDC50A) for protein stabilization (20). In contrast, in our study we observe that loss of CDC50B does not impact on expression of GCα, since GCα-mediated egress still occurs.

The revelation that CDC50C is essential for intraerythrocytic maturation of asexual blood stage *P. falciparum* trophozoites and that CDC50C binds to ATP2 suggests that CDC50C plays a role critical for ATP2 function. In contrast to our findings using native parasite-derived protein preparations, a recent *in-vitro* study using recombinant protein indicated that ATP2 can bind CDC50B; however, CDC50C binding was not tested in that work as the authors could not express it (48). Our study indicates that the essential function of CDC50C cannot be complemented by CDC50B.

Global transposon mutagenesis data suggest that the gene encoding ATP2 is essential in *P. falciparum* blood stages (49) and its orthologue is refractory to targeted deletion in *P. berghei* (50). Whilst its cellular function is unknown, ATP2 has been implicated in resistance to two Medicines for Malaria Venture (MMV) ‘Malaria box’ compounds, mediated through a novel pathway involving gene copy number amplification. Functional characterisation of the mechanism by which drug resistance is achieved remains lacking (51). Interestingly, Cowell et al. observed non-synonymous mutations in genes encoding putative parasite Sec24 and Yip1 proteins (classically involved in vesicular trafficking) in drug resistant parasite lines containing *ATP2* copy number variations (51). Here we found that the ATP2-CDC50C complex influences endocytosis of haemoglobin during blood stage development possibly by influencing the phospholipid makeup of the cytostome, a structure that is crucial for hemoglobin uptake (52, 53), and it remains possible that other endocytic pathways may also be affected by loss of ATP2 function, although these were not investigated. In yeast, different P4-ATPases contribute to distinct vesicular trafficking pathways (43, 44). It is plausible that this could be similar in *P. falciparum*. We speculate that copy number modulation of *ATP2* acquired during selection for drug resistance may modulate the endocytic pathway of the parasite so as to affect drug uptake, although further work is required to investigate this.

A recent study in *P. yoelii* has shown that the orthologue of CDC50C binds to a different P4- ATPase (ATP7) in ookinetes during parasite development within the mosquito (54). This indicates that CDC50C chaperones the activity of distinct P4-ATPases in different developmental stages of the parasite life cycle, in both mammalian and insect hosts. Consistent with this, the transcriptomic profiles of *ATP2* and *ATP7* show that they are confined to asexual and insect stages respectively. The same study demonstrated that the ATP7-CDC50C complex is required for PC uptake in ookinetes and the authors suggested that this process may be required to allow mosquito midgut cell traversal, as CDC50C null or ATP7 null ookinetes could not achieve this. Intriguingly, alignment of ATP7 and ATP2 primary sequences alongside those of model P4- ATPases revealed that the ‘QQ motif’ involved in defining substrate specificity is replaced by QL and QV respectively (Fig S7). Given the similarity between these amino acid motifs, it is plausible that ATP2 also transports PC. Our finding that NDB-PC uptake is unaffected in CDC50C null trophozoites suggests that either ATP2 transports another phospholipid or that lipid uptake in trophozoites occurs via (multiple) redundant pathways. During the preparation of this manuscript, a recent pre-print has found that the *T. gondii* orthologue of CDC50C, CDC50.4, binds ATP2B an essential P4-ATPase that transports PS (55). This CDC50.4-ATP2B complex is required for efficient mirconeme secretion of tachyzoites with no defect observed during parasite intracellular development (55). It is plausible the *P. falciparum* CDC50C-ATP2 complex may perform a similar role in egressed merozoites, however, this was not addressed in our study due to the block of intra-erythrocyte development in CDC50C null parasites. Our work provides substantial new insights into the multifaceted, essential roles played by CDC50C proteins in malaria parasites and highlights potential species-specific divergences in the role of CDC50s in Apicomplexa.

## Materials and methods

### *P. falciparum* culture and synchronisation

*P. falciparum* erythrocytic stages were cultured in human erythrocytes (National Blood Transfusion Service, UK) and RPMI 1640 medium (Life Technologies) supplemented with 0.5% Albumax type II (Gibco), 50 μM hypoxanthine, and 2 mM L-glutamine. Synchronous parasite cultures were obtained as described previously (56). Briefly, late segmented schizonts were enriched by centrifugation on a 60% Percoll (GE Healthcare) cushion, followed by the addition of fresh erythrocytes to allow invasion for 1–2 h under continuously shaking conditions. Remaining schizonts were then removed by sorbitol treatment to yield highly synchronous ring-stage cultures. In all cases, induction of DiCre activity when required was by treatment for 2–4 h with 100 nM RAP (Sigma) as described previously (32, 57). Control parasites were treated with vehicle only (1% v/v DMSO).

### Genetic modification of *P. falciparum* parasites

The CDC50A-HA:loxP, CDC50B-HA:loxP and CDC50C-HA:loxP lines were generated from the DiCre-expressing 3D7 (33) *P. falciparum* clone using SLI of a plasmid containing a SERA2loxPint (57) followed by a triple-HA tag and an in frame Thosea asigna virus 2A (T2A) ribosomal skip peptide and NeoR cassette with a downstream loxP and PbDT 3′UTR sequences as described previously. Re-codonised versions of the C-terminal portion of each gene containing the last transmembrane helix were synthesised commercially (IDT) and inserted downstream of the SERA2loxPint and upstream of the 3×HA tag. Sequences as follows: CDC50A – GATTTCTGGCTCATGAACGAAAAGTACAAGAACGCATTAAACATGAACAATGAGAACGGTTACGGTGACGAAAACAGTCACTTCATAGTTTGGATGAAGACTGCAGCTTTGAGTGAATTTAGAAAGAAGTACGCAAAGATTAACGTAGAGGTAAACTTGCCTATTTACGTTAACATAAACAACAACTTCCCAGTCACCAAGTTCAACGGAAAGAAGTTCTTCGTAATCGCAGAGGGTAGTATTTTCATTAACGAGAAGATTCAGTCTCTCGGTATTCTCTATTTGGTTATAGGTATAATTAGTCTAGGTATAGTTGCATGCCTTATTTACAACCAGATGAAGAATCCGAGGATAATTGGATATCACGCTTATATTTACATCTTCTTCTTCTTGG; CDC50B – GATCACATTTACTTTTGGATGGAGCCTGATATTCAGTACGAGCGTTTGCAGGAGAACAAGGAGACTAAC GAGAAATTGCTAGTTTTGCCTCAGACTTTGAAGTACAACCAGGCTGGTAAGGCAATTGAGAATTCTCACTTCATAAACTGGATGATTCCTAGTGCTCTAAACTACATAAAGCGATTGTACGGAAAGTTGTACATTCCATTGAAGTTCCCCTTCTACATCTACATTGAGAACAACTTCAAGATAAACGACACTAAGATAATCGTAATATCTACATCTCAGTACTACATGAGGACCTTCTTGATCGGCTTTATATTCATCATCATAAGTATCATTGCATTGATCTTGTGCATCTTCTACCTCATCAGGATGAACAAGTACGAGAACAAG; CDC50C – GATGAGTGGAACGCTAAGAAAAGTTTCCAGCTTGTGAGTCTTCGTTCTATTGGTAACTCAAGTTTCAAGTTAGCCTACGCATTCTTTCTTTTAAGTTTGTTGTATTTCATCATGATTATATTCATATTGGTTTTGGTGAAGTGCAAGTACTATAAATTGGGTAAGACTCTTACATACTGTAAGTTATCTATGAACAAGAACATTGAGAAGATGAACTCAAGGAAGAAGACTAACATTCAGAACATTAACAAGAAAATAAACAGTATGCAGCTTGAGATAAT GCATAAAGCCTCATCAGATCCTAACAATCTTGCTGGTGCTGACCACAGTCAGAAGTTGTGTTTCTGTCCATTGCATG. An 800-bp 5′ homology region comprising the native gene sequence upstream of the re-codonised region was cloned upstream of the SERA2loxPint. Following transfection of purified schizonts using an AMAXA nucleofector 4D (Lonza) and P3 reagent, modified parasites were selected as described previously (58).

Oligonucleotide primers used in diagnostic PCR to detect integration and excision of transgenes, and the sequences of re-codonised regions, are provided below in Tables 1 and 2.

CDC50B-HA:loxP GCα-mCherry was generated by transfection of CDC05B-HA:loxP. A linearised donor DNA which inserted mCherry in-frame with the C-terminus of GCα followed by a T2A peptide and BSD selection marker when integrated, and three pDC2-based (33) Cas9 gRNA plasmids were co-transfected, each with different sgRNA targeting the C-terminus of GCα.

sgRNA sequences as follows: sgRNA1 CTCTAAATTATTACAAAATA, sgRNA 2 AGAAAAAACATTCAAGTATC, sgRNA 3 ACGATGAAAAAAAGAAGAAG. Parasites were left to grow for two days post transfection followed by treatment with 5 µg/ml BSD to select for integrants. After the emergence of BSD resistant parasites gDNA was screened for correct integration.

Donor sequences were constructed by amplifying a T2A BSD sequence from pDCIn (DiCre induction) (17) by PCR and cloning using a BsrGI site in frame with the C-terminus of a donor DNA targeting GCα which had previously been constructed in the lab (18).

### Parasite sample preparation and western blot

Parasite culture supernatant samples for egress and adhesin shedding assays were prepared from tightly synchronised cultures as previously described ref. Percoll-purified mature schizonts were resuspended in complete medium and allowed to further mature for 3 h until predominantly mature segmented schizonts. The experiment was then initiated by washing parasites with RPMI three times followed by final re-suspension at a 10% haematocrit in fresh warm RPMI medium. Culture supernatant aliquots (100 μL) were harvested at specified time points by centrifugation. The schizont pellet from t = 0 was retained as a pellet control sample.

Parasite extracts were prepared from Percoll-purified schizonts treated with 0.15% w/v saponin to remove erythrocyte material. To solubilise parasite proteins, PBS-washed saponin-treated parasite pellets were resuspended in three volumes of NP-40 extraction buffer (10 mM Tris, 150 mM NaCl, 0.5 mM EDTA, 1% NP40, pH 7.5, with 1× protease inhibitors (Roche). Samples were gently vortexed and incubated on ice for 10 min followed by centrifugation at 12,000g for 10 min at 4°C. For western blot, SDS-solubilised proteins were electrophoresed on 4%-15% Mini- PROTEAN TGX Stain-Free Protein Gels (Bio-Rad) under reducing conditions and proteins transferred to nitrocellulose membranes using a semidry Trans-Blot Turbo Transfer System (Bio-Rad). Antibody reactions were carried out in 1% skimmed milk in PBS with 0.1% Tween-20 and washed in PBS with 0.1% Tween-20. Appropriate horseradish peroxide-conjugated secondary antibodies were used, and antibody-bound washed membranes were incubated with Clarity Western ECL substrate (Bio-Rad) and visualised using a ChemiDoc (Bio-Rad).

Antibodies used for western blots presented in this work were as follows: anti-HA monoclonal antibody (mAb) 3F10 (diluted 1:2,000) (Roche); mouse anti-GAPDH mAb (1:20,000); rabbit anti- SERA5 polyclonal antibody (1:2,000); rabbit anti-mCherry (1:2000) (Abcam); rabbit anti- haemoglobin polyclonal antibody (1:2,000) (Sigma). Densitometry quantifications were performed using ImageJ.

### Immunofluorescence assays

Thin blood films were fixed with 4% formaldehyde in PBS and permeabilised with PBS containing 0.1% (v/v) Triton X-100. Blocking and antibody binding was performed in PBS 3% BSA w/v at room temperature. Slides were mounted with ProLong Gold Antifade Mountant containing DAPI (Thermo Fisher Scientific). Images were acquired with a NIKON Eclipse Ti fluorescence microscope fitted with a Hamamatsu C11440 digital camera and overlaid in ICY bioimage analysis software or Image J. Super-resolution images were acquired using a Zeiss LSM880 confocal microscope with Airyscan detector in Airyscan SR mode. Antibodies used for IFA were as follows: anti-HA monoclonal antibody (mAb) 3F10 (diluted 1:200) (Roche); mouse anti-PMV mAb (1:50); rabbit anti-ERD2 polyclonal antibody (1:2,000); rabbit anti-EXP2 polyclonal antibody (1:500) (Abcam); rabbit anti-mCherry polyclonal antibody (1:200) (Abcam).

### Flow cytometry

For growth assays, synchronous ring-stage parasites were adjusted to a 0.1% parasitaemia 1% haematocrit suspension and dispensed in triplicate into six-well plates. Samples of 100 μL were harvested at days 0, 2, 4 and 6 for each well and fixed with 4% formaldehyde 0.2% glutaraldehyde in PBS. Fixed samples were stained with SYBR green and analysed by flow cytometry.

### Fluorescent lipid labelling

NBD-PC, NBD-PE, NBD-PS (Avanti polar lipids) were dried and re-suspended in RPMI to 1 mM stock solutions and stored at -20°C. Relevant parasite stages (trophozoites or late schizonts) from a highly synchronous cultures were pelleted and washed twice with RPMI. Parasites were then re-suspended in RPMI containing Hoechst with 1 µM of NBD lipid or no lipid (negative control). Suspensions were incubated at 37°C for 30 minutes and subsequently pelleted by centrifugation. Pellets were then washed three times with pre-warmed RPMI containing 5% BSA followed by resuspension in PBS. Suspensions were then diluted 1:10 and analysed by flow cytometry on an Attune NxT. Samples were gated for Hoechst DNA positivity and the resultant population gated for NBD lipid fluorescence. For trophozoite samples a low Hoechst signal population was gated, and for schizont samples a high Hoechst signal population.

### Immuno-precipitation

Tightly synchronised schizonts (∼45 h old) of CDC50B-HA:loxP, CDC50B-HA:loxP GCα-mCherry, CDC50C-HA:loxP and 3D7DiCre parental parasites were enriched on a 70% Percoll cushion. The schizonts were treated for 3 h with 1 μM C2 (to arrest egress) after which the cultures were treated with 0.15% saponin in PBS containing cOmplete Mini EDTA-free Protease and PhosSTOP Phosphatase inhibitor cocktails (both Roche) for 10 min at 4°C to lyse the host erythrocytes.

Samples were washed twice in PBS containing protease and phosphatase inhibitors, snap-frozen and pellets stored at −80°C. Parasite pellets (70-100 μl packed volume) were resuspended in three volumes of NP-40 extraction buffer (10 mM Tris, 150 mM NaCl, 0.5 mM EDTA, 1% NP40, pH 7.5, with 1× protease inhibitors (Roche). Samples were gently vortexed and incubated on ice for 10 min followed by centrifugation at 12,000g for 10 min at 4°C. Clarified lysates were then added to anti-HA antibody conjugated magnetic beads (Thermo Scientific) or RFP trap beads (Chromotek) which had been equilibrated in NP-40 extraction buffer. Samples were incubated at room temperature for 2 h on a rotating wheel after which beads were precipitated using a magnetic sample rack. The supernatant was removed, and beads washed three times with NP-40 extraction buffer followed by three washes with extraction buffer lacking detergent. Washed beads were then resuspended in trypsinisation buffer (50 mM ammonium bicarbonate, 40 mM 2-chloroacetamide and 10 mM Tris-(2-carboxyethyl) phosphine hydrochloride) and samples reduced and alkylated by heated to 70°C for 5 minutes. 250 ng of trypsin was added to the samples and heated at 37°C overnight with gentle agitation followed by filtration using a 0.22 µm Costar® Spin-X® centrifuge tube filter (Sigma). Samples were then run on a LTQ-Orbitrap-Velos mass spectrometer (Thermo Scientific). Search engines, Mascot (http://www.matrixscience.com/) and MaxQuant (https://www.maxquant.org/) were used for mass spectrometry data analysis. The PlasmoDB database was used for protein annotation. Peptide and proteins having minimum threshold of 95% were used for further proteomic analysis and peptide traces analysed using Scaffold4.

### Measurement of hemozoin content

A culture of 5% parasitemia 1 h synchronised rings stage CDC50C parasites were treated at 1 h post invasion with DMSO or RAP (100 nM) and then left to develop until the early trophozoite stage at 36 h post-invasion. Parasites were then harvested by saponin lysis and then processed similarly to a reported method (46) to purify hemozoin. Pellets were then de-polymerised in 0.5 ml of 0.2 M NaOH solution and the resultant heme content measured by absorbance at 410 nm in a Spectramax iD5 plate reader.

### Measurement of intracellular cyclic nucleotide levels

cAMP and cGMP in mature CDC50B schizonts were measured using enzyme-linked immunosorbent assay (ELISA)-based high-sensitivity direct cAMP and cGMP colorimetric assay kits (Enzo Life Sciences). Mature schizonts were Percoll purified from RAP- or DMSO-treated CDC50B-HA:loxP cultures followed by resuspension and lysis in 0.1 M HCl solution. Samples were pelleted at 10,000 × g, and the supernatant was collected and stored at –80°C until required. To perform the ELISA, samples and standards were acetylated to improve detection sensitivity according to the manufacturer’s instructions. Standards and samples were run in triplicate on the same plate and absorbance at 410 nm read with a Spectramax iD5 plate reader. The standard was fitted to a sigmoidal curve and used to determine cyclic nucleotide concentrations in parasite samples. Remaining supernatant was assayed for protein concentration by a Bradford assay kit (Pierce). cGMP and cAMP reading were normalised by protein content from the Bradford assay.

## Acknowledgments

We would like to thank the following for the kind gifts of antibodies used in this work: Daniel Goldberg (Washington University, St. Louis, MO) for the mouse anti-plasmepsin V mAb; Claudia Daubenberger (SwissTPH, Basel, Switzerland) for the mouse anti-GAPDH mAb; Christiaan van Ooij (LSHTM, London, UK) for the rabbit anti-EXP2 polyclonal antibody.

## Author contributions

All experiments were designed and carried out by AP. MJB and DAB supervised the work overall and SDN designed and provided plasmids for CRISPR editing of GCα.

## List of abbreviations

ATP2: phospholipid-transporting ATPase 2
ATP7: ATPase seven
cAMP: cyclic AMP
Cas9: CRISPR associated protein nine
CDC50: cell division control protein 50
cGMP: cyclic GMP
CRISPR: clustered regularly interspersed short palindromic repeats
DiCre: dimerisable Cre-recombinase DMSO - dimethyl sulfoxide
ERD2: endoplasmic reticulum retention defective two
EXP2: exported protein two
GAPDH: glyceraldehyde three phosphate dehydrogenase
GCα: guanylyl cyclase alpha
GCβ: guanylyl cyclase beta
HA3: triple hemagglutinin epitope tag
IFA: immunofluorescence assay
mCherry: monomeric cherry fluorescent protein
MMV: Medicines for Malaria Venture
NBD-PC: nitro-benzoxadiazol phosphatidylcholine
NBD-PE: nitro-benzoxadiazol phosphatidylethanolamine
NBD-PS: nitro-benzoxadiazol phosphatidylserine
P4-ATPase: type IV ATPases
PC: phosphatidylcholine
PE: phosphatidylethanolamine
PMV: plasmepsin five
PS: phosphatidylserine
RAP: rapamycin
RFP: red fluorescent protein
SBP: skeleton binding protein
SERA5: serine repeat antigen 5
sgRNA: single guide RNA
SLI: selection-linked integration

